# An Open-Source Tool for Automated Human-Level Circling Behavior Detection

**DOI:** 10.1101/2023.05.30.540066

**Authors:** O.R. Stanley, A. Swaminathan, E. Wojahn, Z. M. Ahmed, K. E. Cullen

**Affiliations:** Dept. Biomedical Engineering; Johns Hopkins University; Depts. Otorhinolaryngology-Head & Neck Surgery, Biochemistry & Molecular Biology, Ophthalmology; University of Maryland School of Medicine; Depts. Neuroscience, Otolaryngology-Head & Neck Surgery, Johns Hopkins University

## Abstract

Quantifying behavior and relating it to underlying biological states is of paramount importance in many life science fields. Although barriers to recording postural data have been reduced by progress in deep-learning-based computer vision tools for keypoint tracking, extracting specific behaviors from this data remains challenging. Manual behavior coding, the present gold standard, is labor-intensive and subject to intra-and inter-observer variability. Automatic methods are stymied by the difficulty of explicitly defining complex behaviors, even ones which appear obvious to the human eye. Here, we demonstrate an effective technique for detecting one such behavior, a form of locomotion characterized by stereotyped spinning, termed ’circling’. Though circling has an extensive history as a behavioral marker, at present there exists no standard automated detection method. Accordingly, we developed a technique to identify instances of the behavior by applying simple postprocessing to markerless keypoint data from videos of freely-exploring (*Cib2^-/-^;Cib3^-/-^*) mutant mice, a strain we previously found to exhibit circling. Our technique agrees with human consensus at the same level as do individual observers, and it achieves >90% accuracy in discriminating videos of wild type mice from videos of mutants. As using this technique requires no experience writing or modifying code, it also provides a convenient, noninvasive, quantitative tool for analyzing circling mouse models. Additionally, as our approach was agnostic to the underlying behavior, these results support the feasibility of algorithmically detecting specific, research-relevant behaviors using readily-interpretable parameters tuned on the basis of human consensus.

## INTRODUCTION

Observable actions serve as noninvasive readouts of underlying biological facts - e.g. injury, disease, gene expression, or neural function. This recognition sits at the core of behavioral sciences. Behavioral analysis has historically relied, and largely still relies, on labor-intensive manual behavior coding of real-time or videotaped behavior as its gold standard. Unfortunately, especially when analyzing long sessions or large numbers of sessions, manual coding can suffer due to rater variability, fatigue, or quirks in precise definitions^1–3^). Though recent advances in computer vision^4^ and miniaturized sensors^5^, ^6^ have made quantitative data increasingly available, classifying kinematic data into specific behaviors remains a central challenge. Critically, behaviors which seem clear-cut to a human observer can in reality be noisy and subjective. As a result, while algorithmic behavior detection holds the potential for rapid, objective quantification, automated methods thus face difficulty in explicitly defining complex behaviors.

As one example, rodent behavioral neuroscience studies frequently report a repetitive spinning behavior known as ’circling’. The utility of circling as a behavioral marker, and the resultant need for objective quantification of the behavior, has been recognized for more than 50 years^7^. Examples include studies of basal ganglia damage^7–12^) as well as genetically engineered models of Alzheimer’s Disease^13^, ^14^ and autism^15–17^). Additionally, a large number of mutant mouse strains displaying circling behavior display dysfunction of the vestibular system, which in healthy animals contributes to maintaining balance, steadying gaze, and keeping track of the body within the environment. These include mutants that exhibit loss of vestibular hair cells^18^, disrupted development of stereocilia^19-20^, or disrupted structural development of the inner ear^21-22^.

However, despite its long history as a behavioral marker, the existing literature lacks a standard, quantitative definition of circling. Rather, studies often report the simple presence or absence of circling^5,23–25^. Studies which quantify frequency of occurrence rely on manual coding and use disparate definitions such as complete rotations^26-27^, sequences of complete rotations^13^, or 270-degree turns during which the body travels a minimum distance^28^. Older studies which deployed video analysis relied on tracking the center of mass of a mouse against a high-contrast background^9,10^ and could thus apply limited analysis, whereas more recent work has incorporated commercial and open source tracking solutions but focused on total amount of rotation^29,30^ rather than circling *per se*. Accurate and objective quantification of behavioral parameters such as the frequency of occurrence, duration of bouts of circling behavior, and velocity of movements would facilitate comparison between specific etiologies. This inconsistency reduces the utility of circling as a tool for comparisons across models and setups; thus the need for a broadly-accessible, quantitative, automated tool for the detection of circling behavior.

Here, we present a technique for detecting circling behavior by tuning algorithmic parameters based on consensus occurrence times among human observers. Our technique achieves statistical equivalence to individual observers’ independent labels at matching ultimate consensus times. Specifically, we assessed (*Cib2^-/-^;Cib3^-/-^*) dual knockout mice, a mouse mutant strain we and others have previously reported exhibiting circling^31–33,32^. We track the snout and base of the tail in mice using the open-source software package DeepLabCut^4^ ("DLC"), then compare the performance of several behavioral detection algorithms which apply straightforward analyses to characteristics of the animals’ paths. By identifying the behavioral parameters that result in labels most closely matching human behavioral coding, we develop a quantitative definition of circling behavior for use in comparing against other etiologies. Our methodology provides a simple, replicable, and inexpensive process for recording and quantifying mouse circling behavior. Furthermore, the success of our technique suggests its applicability in detecting and quantifying other research-relevant behaviors.

## RESULTS

The goal of the present work was to build and validate a tool to automatically identify mice exhibiting circling behavior from videos of their free exploration. **Figure 1** illustrates an overview of our data collection and analysis pipeline. We first recorded a set of videos of both wild-type and (*Cib2^-/-^;Cib3^-/-^*) dual knockout mice in six different recording conditions (enumerated in *Methods: Data Generation*). To establish a standard against which to compare the quality of automatic circling detection, six behavioral validation videos - one video of a mutant mouse from each recording condition - were pulled out of this dataset. Three human observers (O.R.S., A.S., E.W.) independently generated lists of times at which circling behavior occurred in these videos. These behavior labels were compared and discussed to produce a set of consensus occurrences, which served as our gold standard (see *Methods: Ground Truth*). All other videos were used to train a computer vision model to track the tip of the snout and base of the tail.

**Figure 1.**
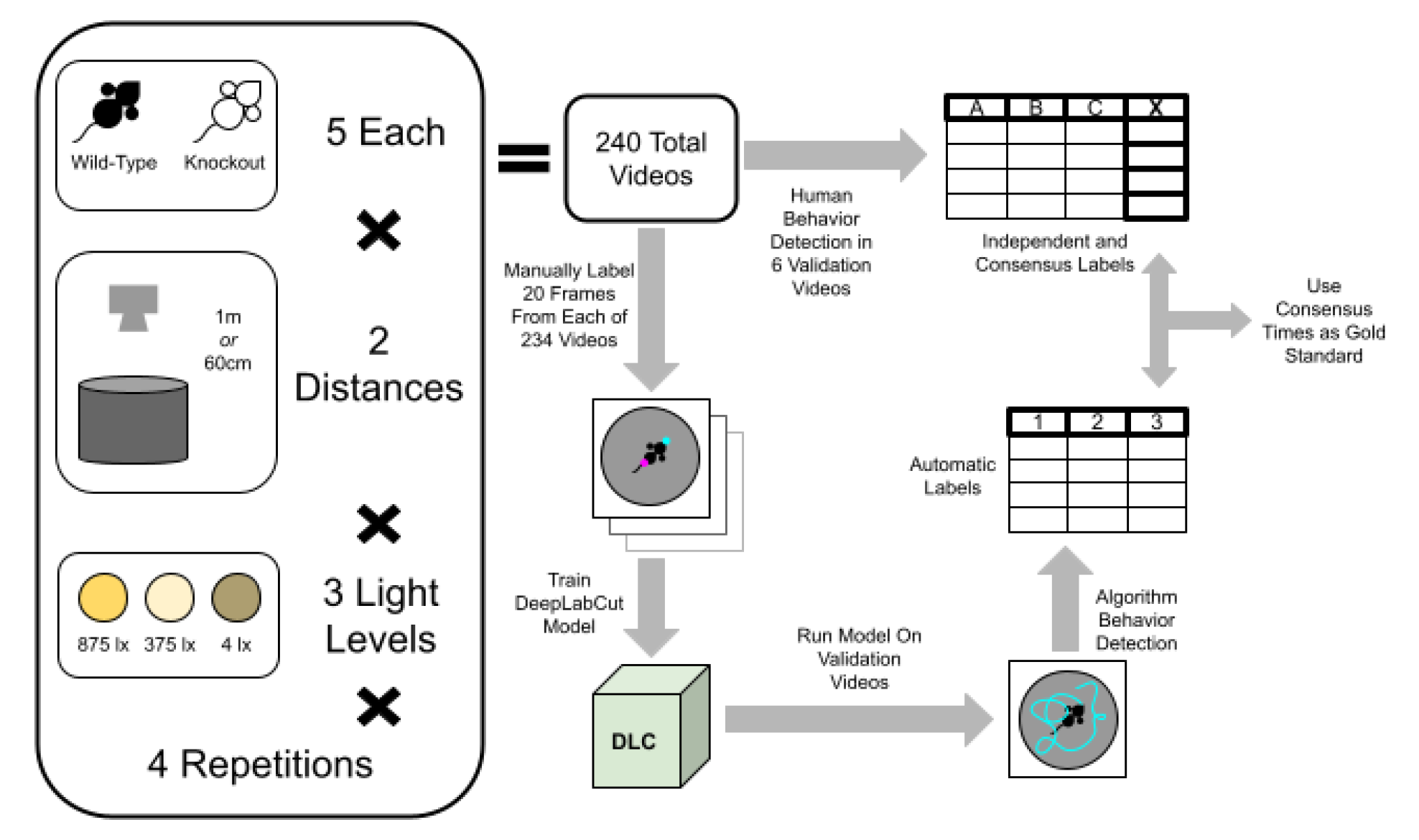
Data Collection Conditions and Analysis Pipeline. We collected videos of five wild-type and five *(Cib2*^-/-^*;Cib3*^-/-^) dual knockout mice exploring a 30cm-diameter cylindrical arena. Each of 6 combinations of light and distance conditions was repeated 4 times for each mouse, resulting in a total of 240 videos. After behavior videos were recorded, 6 videos (one mutant mouse video from each recording condition) were set aside for human behavioral labeling. For each of these held-out videos, three experimenters independently marked occurrences of circling behavior. These behavioral labels were compared and discussed to produce a set of consensus labels. Positions of the snout and tailbase were manually labeled in 20 random frames from each of the remaining 234 videos (4680 total labeled frames). Manually labeled bodypart locations were used to train a computer vision model using DeepLabCut. This trained model was then used to track animals in the 6 held-out validation videos, and the resulting paths were analyzed by three candidate circling detection algorithms. The behavioral occurrence labels produced by these algorithms were compared to human consensus labels to assess performance.

Once trained, the model was run on the behavioral validation videos to generate tracking information, and we applied three progressively more sophisticated algorithms to the resulting keypoint position data. Parameters for each algorithm were optimized to match consensus times as closely as possible as measured by F1 score (see *Methods: Performance Metrics*). Finally, in order to establish the amount of data required to achieve human-level performance at detecting circling behavior, we trained additional computer vision models on subsets of our overall dataset and repeated the above process for only the most effective algorithm.

### Comparison and Consensus Among Human Labelers

To obtain a standard against which to measure our behavioral labeling algorithms, we examined the reliability of and degree of consensus among human labels of circling behavior. To this end, we selected one video of a mutant mouse in each of our six recording conditions at random. Each of these manual behavior validation videos was then viewed and manually screened for instances of circling independently by three experimenters, who were instructed to mark times at which they observed complete rotations during circling behavior but not during exploratory behavior (e.g. turning around after reaching the edge of the arena).

We initially hypothesized that human observers marking occurrences of circling behavior in videos of knockout mice would show strong agreement. To assess agreement among human labelers, F1 scores for each of the 3 possible pairs of independent labels were calculated using one pair member as the ground truth for the other. This resulted in a set of scores from each of three pairs of labels (**Figure 2A**). To our surprise, one pair of labelers showed strong agreement (F1 score of 0.791, 95% confidence interval 0.757 - 0.809, bootstrap) while the other pairs did not (both F1 scores 0.502, 95% CIs 0.4 - 0.563 and 0.415 - 0.549) as a result of one labeler marking substantially fewer instances as true circling behavior. This variability suggested that an automated detection system for circling behavior might not only be more convenient but also more consistent than human observation.

**Figure 2.**
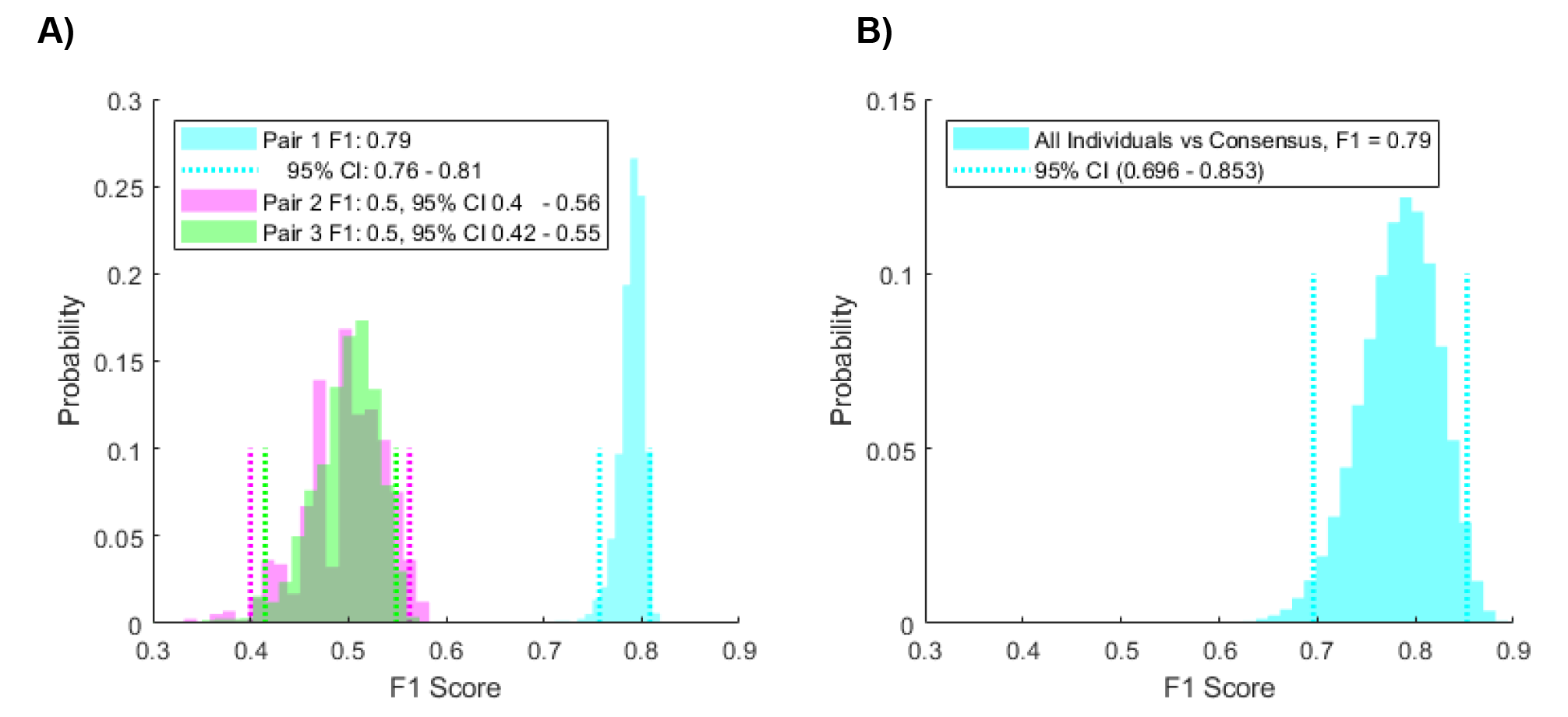
Human F1 Scores. **A)** Using independent observers’ labels as ground-truth for one another demonstrates that human coders vary widely in levels of agreement. For each paired set of human-labeled occurrences, one was treated as ground truth, and the other was scored against it. One pair of labelers showed strong agreement while the other two possible pairings did not. This resulted in F1 scores of 0.79 (95% confidence interval 0.76 - 0.81), 0.5 (95% CI 0.4 - 0.56), and 0.5 (95% CI 0.42 - 0.52). F1 score distributions were generated by summing counts of true-positives, false-positives, and false-negatives for each possible combination (with replacement) of the six validation videos. **B)** Distribution of independent labeler F1 scores versus consensus labels (0.785, 95% CI 0.696 - 0.853). F1 score distribution was generated by bootstrapping 50,000 times from counts of true-positives, false-positives, and false-negatives among the eighteen possible experimenter-video combinations.

We next considered how well independent labels matched the group consensus of circling occurrence times. Times which at least two of the three labelers agreed after group discussion were included in a set of consensus times. This set was used during subsequent scoring of both human and automatic circling detection. To accommodate variations in the precise timing of marked circling occurrences, times within 6 frames (0.1 seconds) of one another were counted as the same instance, a timeframe chosen to cover 95% of the observed variation between independent observers. Scoring independent labels using consensus occurrence times as ground truth produced an overall F1 score of 0.785 (95% CI 0.696 - 0.853; bootstrap, **Figure 2B**). This level of performance served as the baseline for subsequent assessment of automatic methods.

### Developing and Testing Algorithmic Circling Detection Methods

Accurately tracking an animal’s position, or even the position of many body parts, is insufficient on its own to establish the behavior an animal is exhibiting. Rather, this raw data must be processed once collected. To this end, we applied several candidate algorithms for detecting circling behavior using keypoint tracking data. Each algorithm first detects instances in which the DLC-labeled path of the mouse’s snout intersects itself. Importantly, not all intersections are the result of circling - many will be produced by normal exploratory head movements made by the animals. Each algorithm therefore attempts to exclude false positives (i.e., instances incorrectly marked as circling) using one or more features of the animal’s path between the points of collision. Optimal values of these features, as measured by F1 score against human consensus labels, were identified via a grid search, checking each possible parameter combination over a specified range. **Table 1** lists the ranges of these parameters explored via grid search and the final values chosen for each method.

**Table 1.** Method Parameters. Our circling identification algorithms utilize a range of features, values for which were selected via a grid search.

| Method | Parameter | Searched Range | Final Value |
| --- | --- | --- | --- |
| Duration-Only | Min Duration | 0.1 - 0.75 sec | 0.4 sec |
|  | Max Duration | 1 - 1.75 sec | 1.45 sec |
| Swept-Angle | Min Duration | 0.1 - 0.75 sec | 0.35 sec |
|  | Max Duration | 1 - 1.75 sec | 1.45 sec |
|  | Min Consistency | 50 - 70% | 56% |
|  | Min Rotations | 0.025 - 0.75 (9 - 270 deg) | 0.1 (36 degrees) |
| Box-Angle | Min Duration | 0.1 - 0.75 sec | 0.35 sec |
|  | Max Duration | 1 - 1.75 sec | 1.45 sec |
|  | Min Consistency | 50 - 70% | 56% |
|  | Min Rotations | 0.025 - 0.75 (9 - 270 deg) | 0.1 (36 degrees) |
|  | Min Size | 0.1 - 0.7x body length | 0.5x |
|  | Max Aspect Ratio | 2.0 - 2.75 | 2.5 |

#### Duration-Only Method

In order to establish a lower bound on automatic behavioral detection performance, we first assessed a simple detection algorithm which considers only how long a candidate circle takes. The ’Duration-Only’ method locates points at which the path of the mouse’s snout crosses over itself and excludes those which are either too short or too long (**Figure 3B, Row 1**). Our parameter grid search produced an F1 score of 0.66 (95% CI: 0.56 - 0.73, calculated across all validation video combinations) using duration boundaries of 0.4 to 1.45 seconds (**Figure 3C, Row 1**). The performance level achieved by even this straightforward method is not statistically significantly lower than human performance, though only marginally (p = 0.0665, exact two-tailed Wilcoxson signed rank test of automatic scores versus human scores). We noted that, as expected given its simplicity, this method was not sufficient to filter out many false positives. Specifically, it incorrectly labeled many cases of head-only exploratory movements as circling, some of which are illustrated in **Figure 3D, Row 1**. To filter these out more effectively and thereby improve behavioral detection, we next explored incorporating the labeled tailbase position.

**Figure 3.**
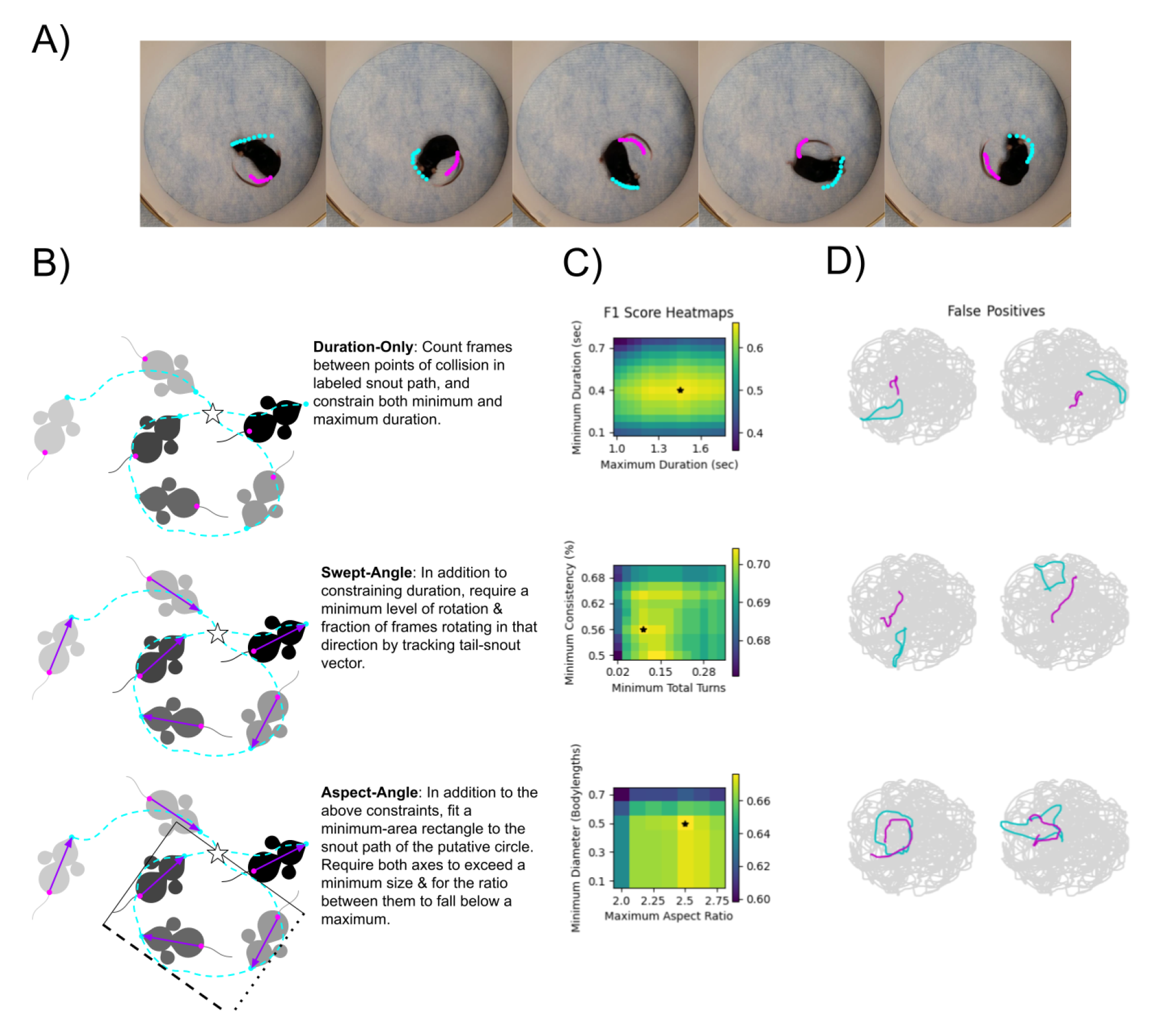
Method Parameters & Performance Levels. **A)** Example DLC-labeled frames, showing trailing labeled points. **B)** Illustration of circle detection using each of the described methods. Duration-Only considers only time taken to complete the circle, Swept-Angle additionally calculates the angle of the tail-to-snout vector for each frame and considers both its total movement and its ’consistency’ (fraction of time moving in the predominant direction), and Aspect-Angle constraints the geometry of the circle based on the aspect ratio and minor axis length of the circle’s minimum bounding rectangle. **C)** Heatmaps of F1 scores when varying two of each methods’ two parameters. Maximum F1 scores are indicated by stars; 0.66 (min duration 0.4 sec, max duration 1.45), 0.7 (0.35 - 1.45 second duration, 56% min consistency, min rotation of 36 degrees), and 0.74 (same best parameters as Swept-Angle, plus min axis size of 0.5 bodylengths and max aspect ratio of 2.5), respectively. **D)** Examples of false-positive detection using each method in one validation video.

#### Swept-Angle Method

For a mouse that is spinning rather than exploring with its snout alone, the angle of its snout relative to its tail should change noticeably and in a consistent direction. Accordingly, we next considered what we term the ’Swept-Angle’ method for excluding false-positive instances of circling. As illustrated in **Figure 3B, Row 2**, this technique calculates the angle of the mouse’s body in each frame using the vector from labeled tailbase position to labeled snout position. It then screens candidate circles using bounds on duration as well as minimum total rotation and minimum rotational consistency. Total angular change was measured by simply summing frame-by-frame changes, while the consistency of rotation was calculated as the fraction of the time the body vector changed angle in its more prevalent direction (i.e., clockwise vs. counterclockwise).

We performed a grid search over the four parameters of minimum duration, maximum duration, minimum rotation, and minimum turning consistency. The Swept-Angle method reached an F1 score of 0.7 (95% CI 0.6 - 0.78, calculated across all validation video combinations) using a minimum time of 0.35 seconds, a maximum time of 1.45 seconds, minimum circling consistency of 63%, and a minimum net rotation of 36 degrees (0.1 full rotations); see sample heatmap, **Figure 3C, Row 2**. This statistically matches independent human performance (p = 0.495, exact two-tailed Wilcoxson signed rank test of automatic scores versus human scores on each validation video), and it outperforms the Duration-Only method (p=0.0312). Notably, this means that the Swept-Angle method outperforms the Duration-Only method more consistently than independent human scoring, though it does not do so by as large a margin on average. Thus, incorporating the additional information of the tail’s position resulted in a modest but detectable increase in behavioral detection performance. In examining erroneous circles detected by this method (see examples plotted in **Figure 4D, Row 2**), we identified many cases in which false positives were oblong. To counteract this, we next attempted to incorporate additional geometric information about candidate circles.

**Figure 4.**
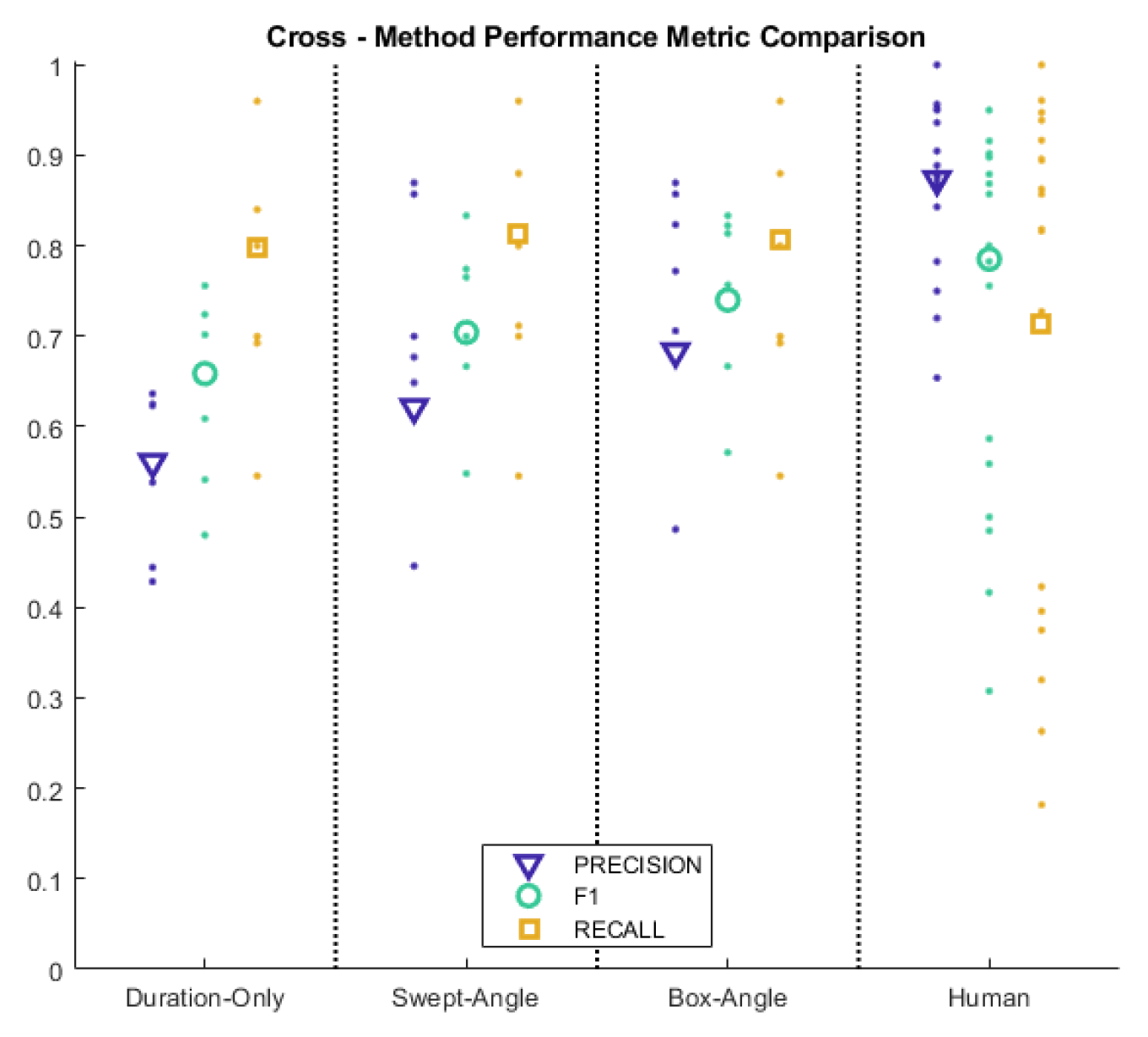
Cross-Method Performance Comparison. Each column contains weighted-average performance metrics (precision: blue triangles, left; F1 score: teal circles, center; recall, yellow squares, right) across the validation video set using the Full Dataset model. Large markers represent overall scores while small dots represent individual videos. Notably, in contrast to the results from independent human labels, each automatic detection method produces higher recall than precision.

#### Box-Angle Method

To filter out cases in which the animal’s path is insufficiently circular, we implemented a final method, which contains an additional step in which a candidate circle is fitted to a minimum-area rectangle to provide additional geometric information (see **Figure 3B, Row 3**). In addition to the metrics described for the Swept-Angle method above, the resulting ’Box-Angle’ method constrains the minimum side length and maximum aspect ratio of the resulting fitted rectangle. To avoid relying on specific information about the camera and the geometry of the recording setup, the minimum side length is specified relative to the median size of the mouse as measured by tail-to-snout vector over the course of the video. Our parameter grid search (see heatmap, **Figure 3C, Row 3**) resulted in duration limits of 0.35-1.45 seconds, 56% minimum consistency, 36 degree minimum rotation, 0.5x body length minimum diameter, and a maximum aspect ratio of 2.5; this combination produced an F1 score of 0.74 (95% CI 0.632 - 0.818, calculated across all validation video combinations). This performance level again statistically matches human performance (p = 0.61, exact two-tailed Wilcoxson signed rank test of automatic scores versus human scores), while outperforming the Duration-Only method (p=0.0312; exact two-tailed Wilcoxson signed rank test of the 6 video scores from each method). While it successfully avoids some of the oblong false-positive circles observed using the Swept-Angle method (**Figure 3D, Row 3**), the difference in performance does not reach significance (p=0.125).

### Comparison Across Methods

While these methods varied in their precise levels of performance and the variability of that performance, incorporating additional steps to filter out false positives generally increased an algorithm’s overall F1 score. Notably, and in contrast to the trend observed in independent human labels scored against human consensus, all the automatic detection methods show greater recall than precision (see **Figure 4**). Based on the fact that it performed at least as well on each validation video as the Swept-Angle method, we chose to use the Box-Angle method for subsequent comparisons among DLC models trained on different subsets of the 4680 frames used for the Full Data model.

### Performance Among Datasets

All the results reported thus far were produced using a DLC-trained convolutional neural network model, termed the "Full Dataset" model, which used 20 manually-labeled frames from each of 234 mouse behavior videos not used for human labeling validation. This model represents a substantial investment of experimenter effort, raising the question of whether sufficient behavioral detection performance could be achieved more easily.In particular, DLC’s creators observed good keypoint labeling performance with substantially smaller datasets^4^, but it was unclear *a priori* what labeling quality is necessary for successful behavior detection.

In order to investigate the minimum amount of data and thus human labor needed to obtain good automatic behavioral detection performance, as well as to examine the relative importance of various aspects of dataset diversity, we trained several DLC models using different subsets of our manually-labeled frames, detailed in **Table 2**. The ability of each model to accurately label the locations of the snout and tail-base was evaluated using frames that model had not previously seen; as these were not used during training, they provide a measure of each neural network’s ability to accurately label snout and tailbase locations when generalizing to novel videos. Models with fewer than five total videos were all tested on frames from videos of mutant mice in the medium-light condition for the sake of direct comparability. In order to compare all models against the same number of test frames, a randomly selected 15% of frames the available 4680 were held out for out-of-sample testing to determine the model’s ability to generalize to unseen images. Notably, the performance of the Full Dataset model, as measured by mean root-mean-squared error (in pixels), was about half that of models trained on only four videos.

**Table 2.** Keypoint Labeling Performance Among Models. Datasets comprising subsets of our manually-labeled frames were used to train different neural network models using DeepLabCut. All models were initialized using the pretrained ResNet50 model available through DLC and trained for up to 100,000 iterations at a learning rate of 1×10^-3^. Root-mean-squared error in units of pixels between model-assigned and manually-labeled snout and tailbase positions, was calculated on ’test’ frames, i.e. those not directly used to train the neural network. The Single Video model and the models trained on 4 videos were each tested on frames from videos of mutant mice in the medium-light condition for direct comparability. The Full Data model was tested on a randomly held-out 10% of labeled data.

| Network | Single Video | More Animals | More Conditions | Full Data |
| --- | --- | --- | --- | --- |
| Mice | 1 KO | 4 KO | 1 KO | 5 WT<br>5 KO |
| Videos per Condition | 1 | 1 | 1 | 4* |
| Conditions | Near/Bright | Near/Bright | Near/Bright<br>Near/Dark<br>Far/Bright<br>Far/Dark | All |
| # Videos | 1 | 4 | 4 | 234 |
| RMSE (Pixels) | 43.66 | 23 | 22.41 | 8.13 |

\*: Except for a single KO-mouse video from each condition, held out for behavioral validation

To determine whether this difference in pixel-wise error impacted the ability of networks to successfully identify circling behavior, we ran F1-score-optimizing parameter grid searches over the validation videos for each network using the Box-Angle method. As described above, the Full Dataset network achieved an F1 score across the validation videos of approximately 0.74. The Single Video network, trained on the smallest dataset (20 frames from a single video of a mutant mouse) achieved an overall F1 score of 0.659. Models trained on either a single video from each of four different animals (’Multi-Animal’ model) or on the same animal but in different recording conditions (’Multi-Condition’ model) obtained F1 scores of 0.728 and 0.734, respectively (**Figure 5A**). While the Full Dataset model’s performance matched that of independent human labelers (p=0.61; exact two-tailed Wilcoxson signed rank test of automatic scores paired against summed human true positive, false positive, and false negative counts for each video validation video), the Single-Video did so only barely (p = 0.102). The Multi-Animal and Multi-Condition models produced noticeably closer matches to human performance (p = 0.420, and 0.3271, respectively) (**Figure 5B**).Taken together, this analysis suggests human-level performance in detecting circling behavior can be obtained with a surprisingly small investment of experimenter time and effort.

**Figure 5.**
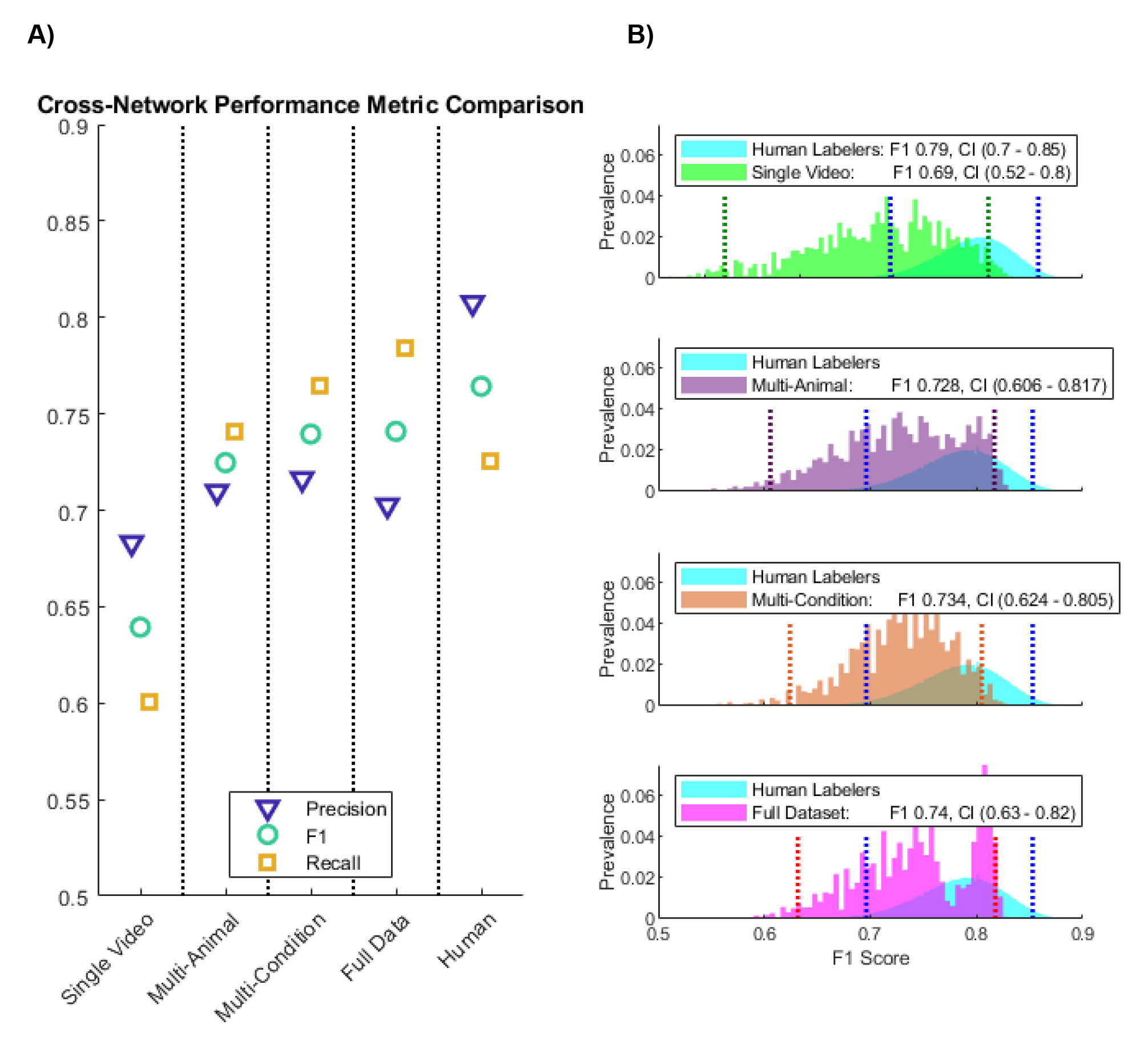
Comparison of Performance Metrics Across Networks. **A)** Performance metrics for each trained network, using individually-optimized Box-Angle method parameters, alongside performance metrics for independent human screening. Each column includes precision (left, blue triangles), F1 score (center, teal circles), and recall (right, yellow squares). **B)** Bootstrapped F1 scores for independent human labeling (cyan) versus automatic labeling via Box-Angle method by each model, all scored against human consensus labels: Single Video model (green), Multi-Animal model (purple), and Multi-Condition model (orange), and Full Dataset model (magenta). Dotted vertical lines represent bounds of 95% confidence intervals.

### Differentiation Between Mutant and Wild-Type Mice

The present work was aimed to develop a tool for detecting vestibular-mutant mice. To assess its success, we therefore applied the final method developed above to all 240 videos collected for training our computer vision models and found the number of putative circling instances identified in each. We then calculated the F1 score obtained by a range of possible thresholds for observed circling instances per-minute by counting mutant videos exceeding the threshold (true positives), mutant videos below the threshold (false negatives), and wild-type videos exceeding the threshold (false positives). The number of average circles per minute observed in wild-type vs. mutant mouse videos differed dramatically (<0.1 vs >50, **Figure 6A**). Using a simple threshold-based classification method produced an F1 score of ∼0.9 at a threshold of 0.95 circles/minute. (**Figure 6B**)

**Figure 6.**
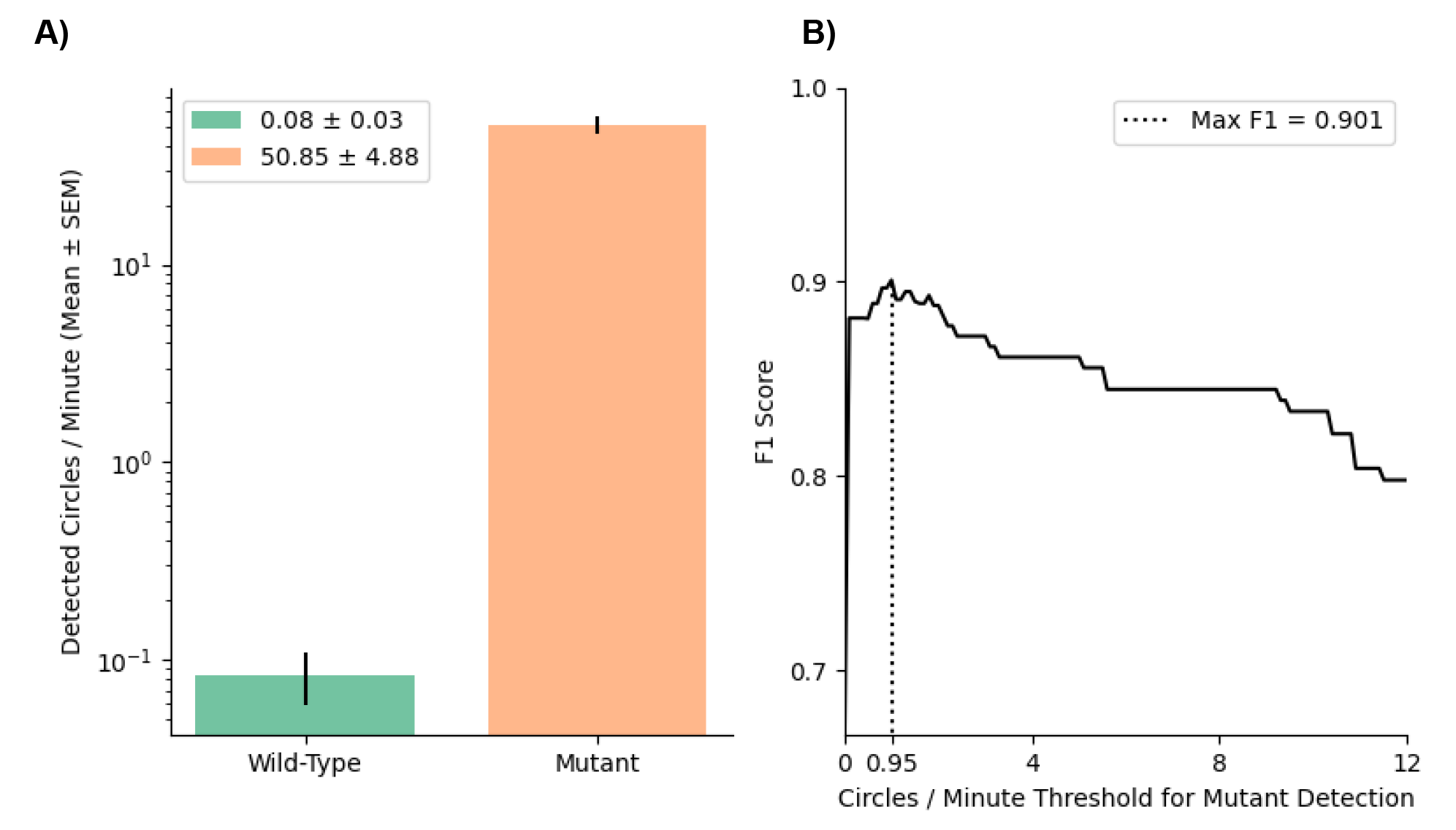
Mouse classification performance metrics. **A)** Videos of wild-type mice contained on average less than one circling instance per ten minutes of footage (green) while (*Cib2^-/-^;Cib3^-/-^*) mutant videos contained on average more than fifty instances of circling per minute (orange). **B)** F1 score for our final algorithm when discriminating between dual knockout and wildtype mouse videos, as a function of circling frequency. Using a simple threshold classifying any video with at least 0.95 circling instances per minute as a mutant (dotted line) produced an F1 score of ∼0.9. Notably, setting any requirement for minimum circling per minute substantially outperforms chance (chance level = 0.66, floor of y-axis).

## DISCUSSION

In the present study, we developed a technique to automatically detect circling behavior in videos of freely exploring mice using readily-available, consumer-grade hardware and open-source software. Importantly, this method is applicable to analyzing information captured by any method for keypoint tracking, whether open-source or proprietary. This makes it a convenient, quantitative tool to screen mice for circling behavior according to specific, objective criteria. More generally, our results suggest that similar procedures to develop consensus behavioral labels among human observers could be straightforwardly applied to enable effective, automatic detection of other behaviors of interest. The development of such quantitative tools with low barriers to entry is essential for the comparative analysis of behavior, as it expands the number of research groups able to produce directly inter-comparable data.

### Limitations of Prior Approaches

Studies which report circling have varied widely in methods of detection and analysis, ranging from qualitatively reporting the presence or absence of circling^5,23–25^ to counting rotations per minute using manual or automated video tracking^9,10,29,30^. Critically, existing methods using video analysis face limitations in quantifying circling behavior due to false positives arising from grooming or exploratory turns. Our work specifically addresses this issue by incorporating filters for excluding false positives using carefully tuned geometric parameters. By tracking only two key points on the mouse and applying straightforward algorithms, we achieved behavioral labeling performance (F1 score ∼0.74) statistically similar to that of independent human coders (∼0.78, p=0.610, **Figure 5B**).

Software to track rodents in videos is commercially available, but such products face issues of both price and opacity; as closed-source software, they limit the ability of researchers to examine how specific results were generated and to customize or otherwise modify those underlying methods.^34^ In the present study, we chose to use the open-source markerless feature-tracking toolbox DeepLabCut ("DLC") to track the positions of keypoints on animals. Notably however, as discussed below, the analysis applied to this positional information is agnostic to the tracking method used.

### Availability & Ease of Use

Our Full-Data model, the Python script to detect circles from keypoint positions, and an associated Anaconda environment file can be found at our GitHub (https://github.com/CullenLab/CirclingDetection) along with a step-by-step guide to installing and using the system. To use our code, we recommend installing Anaconda (https://www.anaconda.com/products/distribution) and using it to create a Python environment from the `.yaml` file included in the GitHub Repository. Once this is done, the code can either be run from the command line from within the downloaded Repository or run via one of the development environments Anaconda offers.

Although we chose to develop our methodology using DLC, the detection algorithm can be employed using any technique to track the snout and tail base. Those utilizing other tracking methods should note that while the `Circling_AspectAngle.py` script assumes a particular configuration of columns in the `.csv` files to be analyzed (described further in the repository documentation), CSVs produced by other tracking methods can be straightforwardly modified to fit this scheme.

Based on performance levels among models trained on different datasets, we found that human-level performance in behavioral detection could be achieved even with small investments of human effort devoted to manual image labeling, which comports with observations by DLC’s developers^4^. Specifically, we found that datasets comprising only 80 manually labeled frames produced performance comparable to that of independent human labelers (**Figure 5B**). We conclude that the tradeoff between time (i.e., required number of labeled images) and accuracy is minimal; close to maximal detection performance can be reached by labeling a relatively small number of frames of high-quality behavioral videos.

### Broader Applicability of this Approach

Recent developments in mouse genetic engineering, involving the creation of transgenic and knockout mutant mice, have provided a novel opportunity to study the relationship between genes and behavior.^3^ For example, many mutant mice strains that have been characterized with an underlying impairment of peripheral vestibular function display a circling behavioral phenotype. While there exist a number of non-invasive methods to detect the existence of vestibular dysfunction in mice, such as the rota-rod and balance beam tests^6^, the variety of vestibular-loss circling mouse models suggests screening for circling may serve as a convenient screening tool for identifying novel models of vestibular dysfunction. In the present study, we specifically assessed the circling behavior (*Cib2^-/-^;Cib3^-/-^*) dual knockout mice^31–33^). CIB2 is found in the stereocilia tips of the receptor cells within the vestibular sensory organs^35^ (i.e., vestibular hair cells), suggesting the circling behavior observed in these mice results specifically from deficits in peripheral mechanotransduction. Notably, the strain studied in the present work is just one example of a large number of strains with mutations homologous to subtypes of Usher syndrome^36^, including deaf circler^26^, waltzer^29^, Ames waltzer^37^, and Jackson shaker^38^ mice. Our results are thus likely to apply relatively directly to this extended web of mouse models. In particular, a method for automatically & objectively quantifying circling behavior may reveal otherwise undetectable differences in circling parameters (e.g. differences in frequency, rotational velocity, and geometry) between model strains.

More broadly, tools for detecting and quantifying behavior which do not rely on specialized experience with computer vision or writing software will allow a wider array of research groups to directly compare analyses of neurodevelopmental differences and intervention effects. The use of shared automatic behavioral analysis tools would increase both the speed and consistency of behavioral labeling, especially when analyzing long or numerous videos. For example, the technique presented in this paper would aid in screening novel mutants for vestibular dysfunction. Common behavior definitions may also provide inter-comparability among studies within and across research groups as well as improving the reproducibility of those studies.

In this context, it is noteworthy that the concepts underlying our approach can be readily applied to other behaviors of interest to researchers. Specifically, by first creating a set of occurrence times coded independently and then collaborating on a set of consensus occurrence times, we were able to directly quantify human-level performance. In principle, this enables working with a wide range of behaviors which may be difficult to define explicitly ahead of time but which we ’know when we see them’. For example, a similar strategy using keypoint tracking to develop computational heuristics has been applied in the identification of freezing behavior^39^, which must be distinguished from simply remaining still, just as circling must be distinguished from normal exploration. In such contexts, our consensus-development step is especially important to avoid incorporating biases or quirks of any one observer.

Furthermore, an important advantage of video-based approaches is that they are non-invasive. This stands in contrast with the emerging use of head-mounted MEMs-based sensors to assess motor impairments in mouse models (see for example ^5^, ^6^), which typically require an experimental surgery to securely fix the sensor to the mouse’s skull. We speculate that as the spatio-temporal resolution of readily available, consumer-grade hardware continues to improve, the tradeoff in resolution between a completely noninvasive recording technique (video analysis) and a more invasive technique in which inertial sensors are mounted surgically will become less critical. Finally, it is noteworthy that incorporating a larger number of keypoint labels will be critical for examining rodent posture and gait^40^, and has been incorporated into other analyses of circling mice specifically^41^. Thus, while the current study focuses on creating as simple a method as possible for detecting a specific behavior, more complex analysis will be required to elucidate subtle differences in the effects of genetic, pharmacological, or electrophysiological interventions on other motor behaviors.

### Conclusions and Implications

Emerging video-based technologies that facilitate the tracking of key body features are well-suited to the development of accessible methods for objectively quantifying behavior, including in comparing normal versus mutant mice. Mouse models are advantageous for studies of treatments and causes of vestibular impairment, due in part to the ability of researchers to genetically engineer new strains using increasingly sophisticated tools.^42^, ^43^ These studies are clinically relevant in light of aging populations, as vestibular dysfunction substantially increases fall risk and causes symptoms including dizziness, vertigo, nausea, and blurred vision. In adults over 40, its prevalence has been estimated as ranging from more than one in twenty^44^ (using vestibular-specific clinical measures) to more than one in three people over 40^45^ (using broader balance-related symptoms). Here, we present an open-source tool which accurately distinguishes between wild-type mice and circling mutant mice. By demonstrating the effectiveness of tuning behavioral detection algorithms based on consensus among multiple human observers, our results not only provide a convenient, quantitative screening method for mouse models of vestibular dysfunction, they also serve as an important step toward standardized, automatic measurement of motor dysfunction in mouse models that provide more reliable measurements across studies and laboratories.

## METHODS

### Animal Care and Housing

Five adult wild-type (C57BL/6) mice (Jackson Laboratories) and five adult mutant (*Cib2-/-;Cib3-/-*) mice were used in this study. Generation of mutant mice is described in a separate paper^33^. Animals were group-housed with their littermates on a 12:12 h light: dark cycle at 20°C with *ad libitum* access to food and water. All animal procedures complied with the National Institutes of Health Guide for the Care and Use of Laboratory Animals and were approved by the Institutional Animal Care and Use Committees (IACUCs) at University of Maryland (protocol #0420002).

### Data Generation

We recorded videos of 5 wild-type mice and 5 (*Cib2^-/-^;Cib3^-/-^*) dual knockout circling mice during single-animal exploration of a cylindrical arena 30cm in diameter, similar to ones used in past studies.^8,9,15^ For each mouse, we recorded four 2-minute videos at 60fps in each of six recording conditions - low (∼4 lux), moderate (∼375 lux), or bright lighting (∼875 lux), each with the camera either near to (60cm) or far from (100cm) the animal. These were selected to span a broad range of potential experimental setups and to provide varied data for neural network training; although we made use of the DLC toolbox’s options for data augmentation via random image perturbation (see *Methods: Neural Network Training*), recording in varied distinct conditions allowed us to examine the effects of qualitative differences in training datasets (i.e., what data is included) rather than only quantitative differences (i.e., how much data is included). Conditions are enumerated in **Figure 1**. Our total set of 240 videos thus consisted of 120 videos of wild-type mice and 120 videos of mutant mice.

### Ground Truth

Assessing the effectiveness of our tool required a ground truth against which we could compare the automatic detection of circling behavior. One mutant mouse video from each recording condition was set aside at random for human behavioral labeling. Because assigning no circles to a WT video would artificially inflate the performance metrics reported here, we excluded videos of WT mice from our validation set.

Consensus behavior labels were established by first having three experimenters independently note occurrences of circling in these videos, then by comparing any discrepancies frame-by-frame as a group. After debating discrepancies, only circling instances on which two of the three experimenters agreed were included in the consensus set. Subsequently, circling instances detected by either human observers or automatic methods were counted as true positives if they fell within 0.1 seconds of a consensus-labeled occurrence. We chose this window to accommodate 95% of discrepancies among independent human labels of the same instance.

### Performance Metrics

To balance the need to avoid both false positive and false negative errors, we used F1 score to assess three methods of detecting circling behavior in DLC-labeled paths, calculated as follows:

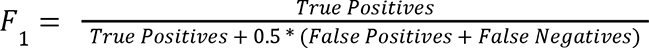

This metric is equivalent to taking the harmonic mean of precision (fraction of all identified circles that are correct) and recall (fraction of true circles identified):

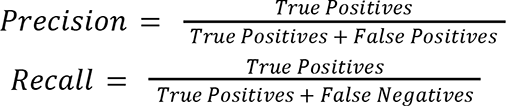

### Algorithm Specifications and Parameter Grid Search

We compared three algorithms for detecting circling using labeled locations of two keypoints on the body of freely exploring mice. All three initially search the path of the mouse’s snout for cases where it crosses over itself between a minimum and maximum duration. After identifying these potential cases of circling behavior, two of the methods consider one or more additional features of the mouse’s path to filter out false positives.

The ’Duration-Only’ method uses only a minimum and maximum duration, the ’Swept-Angle’ method excludes candidate circles based on the magnitude and consistency of change in the angle of the vector from the labeled tailbase position to labeled nose position, and the ’Box-Angle’ method additionally considers the size and aspect ratio of the area swept by the snout during a putative circle. We thus needed to optimize either 2, 4, or 6 parameters depending on the method being considered. This was accomplished via grid search, in which a space of parameters is explored exhaustively. Parameter ranges explored during our grid search, as well as the resulting F1-optimal parameters, are summarized in **Table 1**.

### Neural Network Training

We chose to use DLC, an open-source tool for training deep convolutional neural networks to recognize user-labeled image features, to track the locations of mouse-body keypoints due to its accessibility, as it can be straightforwardly used by researchers with little machine learning experience with consumer-grade computing hardware.

From each of the 234 videos not used in manual behavior validation (120 WT, 114 dual KO), we labeled 20 random frames with the positions of the mouse’s snout and the base of its tail. These manually-placed keypoints served as training data (4680 training frames total). We utilized data augmentation in the form of the ’*imgaug*’ dataloader included in DLC, which applies perturbations during network training such as cropping, blurring, and rotating training images. We refer to this as the "Full Dataset’’ model, in contrast to models trained on subsets of this data.

To determine the number and kinds of videos that were necessary to reach a plateau in performance, we used different subsets of our 234-video dataset to train several different DLC models. To establish a lower bound on the range of DLC model performance, we trained one model on data from a single video of a mutant mouse in a single recording condition (specifically, in bright light and with the camera close to the cylinder; the "Single Mutant Video" model). Then, to assess the impact of dataset size and diversity, additional DLC models were trained on 4-video subsets. (**Table 2**)

Each model was initialized using a 50-layer pretrained network model (ImageNet-trained Resnet50) and trained for 100,000 iterations at a learning rate of 0.001. After training, each DLC network was run on the 6 held-out validation videos to produce position traces to be analyzed using the Box-Angle method described above.

### Statistical Analyses

P-values for differences between F1 score distributions were calculated using exact two-tailed Wilcoxson signed rank tests, in which F1 scores from each method on each video are paired against one another.

Confidence intervals were calculated via bootstrap. Specifically, bootstrapped datasets for each method were obtained by summing the true-positive, false-positive, and false-negative counts from subsets of validation videos resampled with replacement from the data to create a distribution of possible datasets of the same size as the original (i.e. drawing from 1 to N with replacement N times; for independent human performance, we sampled among the N=18 score sets (3 experimenters x 6 videos); all other cases sampled from N=6 videos). For all methods of N=6, all possible combinations were checked (6^6^ = 46,656 combinations). For the sets of independent human labels, exhaustive testing was infeasible (18^18^ ∼ 3.9E22 combinations), so 50,000 random samples were computed. This method was selected over directly bootstrapping and averaging video F1 scores in order to give equal weight to each instance of correctly identified, falsely detected, and missed circling regardless of the video in which it occurred.

## ACKNOWLEDGEMENTS

The authors wish to thank Celia Fernandez-Brillet, Olivia Leavitt, Robyn Mildren, Kiara Quinn, and Kantapon Pum Wiboonsaksakul for their insightful feedback on the manuscript, Dale Roberts for assistance with Python, and Ryan Riegel for assistance with recording behavior videos. This work was supported by NIH grants R01DC012564 and 2T32DC000023.

